# Strain-dependent selective antimicrobial action of cationic polyelectrolytes on Gram-negative bacteria

**DOI:** 10.1101/2021.08.31.458386

**Authors:** Iaroslav Rybkin, Janja Mirtič, Hana Majaron, Jitka Hreščak, Miran Čeh, Janez Štrancar, Julijana Kristl, Tomaž Rijavec, Aleš Lapanje

## Abstract

Although electrostatic modification of bacterial surfaces using polyelectrolytes (PEs) is a convenient and versatile tool for biotechnological processes, the ambiguities in toxicity of PEs between various bacteria and the insufficient understanding of the mechanism of action of cationic PEs and their nano-thick shells formed around the bacteria create a bottleneck of the approach. Here, we show how the viability of two bacterial strains, *Escherichia coli* and *Pseudomonas stutzeri*, both from the Gram-negative group differs, when the cells are exposed to cationic PEs under different conditions. Although the cell wall architecture of the strains should be structurally similar, we found that the viability of *E. coli* was not affected by the electrostatic deposition of polyethyleneimine (PEI) or poly(allylamine) hydrochloride (PAH), whereas for *P. stutzeri* the deposition resulted in high death rates. The cells of *E. coli* proved to be suitable templates for Layer-by-Layer (LbL) modification, while in *P. stutzeri* a modified protocol with mild conditions had to be used to ensure the viability of the cells. Super resolution stimulated emission depletion (STED) microscopy allowed us to clearly visualize that after PE deposition onto the surface of the cells, the PEs could penetrate inside the cells of *P. stutzeri*, while forming a capsule around *E. coli* as expected. Therefore, this knowledge will help us select the most appropriate combinations of strains and PEs, for biotechnological processes or biomedical application, preventing unwanted toxicity.

## Introduction

There is a substantial need to find alternative antimicrobial approaches to fight against microorganisms with antimicrobial resistance (AMR), on the one hand, due to a high speed of their dissemination, and on the other hand, due to a dramatic reduction in the number of antibiotics that can eradicate such infections. The highest impact on the health is made by biofilms as bacteria present within these films are unattainable from the action of antibiotics and antimicrobials compounds.^1,2^ Moreover, cells exchange genes containing AMR resistance between members of biofilms, making the biofilm even more resistant for treatment.^3^

One of the approaches to make a surface bactericidal is to coat the surface with positively charged PEs. However, PEs can be used either of two ways to prevent colonization of unwanted microbes (i) by aiding probiotic or beneficial bacteria to attach the cells on surfaces by electrostatic attachment^4–6^ or (ii) to induce a toxic effect that prevents attachment of unwanted microbes.^7,8^ These two approaches are diametrically opposing since one requires the preservation of bacterial viability, although another is based on the PE toxicity to inhibit cell growth. Therefore, we expect tremendous ambiguity in the conditions and reasons for selection of one sort of PEs over others, either for coating or killing cells.

In the case when PEs are used as mild coatings deposited in the LbL manner,^9^ it allows the formation of ultra-thin (10-20 nm in thickness), but extremely strong nano-thick capsules that are permeable to nutrients.^10,11^ LbL nanocapsules consisting of PEs, firstly functionalize the cell’s surface, by introducing positive charges and secondly facilitate cell’s attachment to surfaces. Since the cells stay under physical constraint in the permeable capsules, it also allows primary studies of cellular physiology.^6^ Although bacterial cells have been suitable templates for PEs deposition, the latter has been noted to negatively affect the viability of various strains in several cases.^7–11^

In the case when PEs are used as coatings, inhibiting cell growth,^7,12–14^ they are known to be particularly toxic to bacteria.^15–17^ However, the toxicity of these coatings was ambiguous as the susceptibility of various bacteria to cationic PEs differed from one strain to another, as well as between Gram groups. ^15,16,18–22^ Thus, we hypothesize that toxicity can be dependent either on the structural differences of PE molecules or on the diversity of the structure of the cell envelope. In particular, cationic PEs can have different molecular weights (MWs) or distances between functional groups, their hydrophilic-hydrophobic balance and the nature of counterions, which all can contribute to the observed toxic effects.^12,15,22–24^

On the other hand, we can expect that the architecture of the cell envelope of Gram-negative or Gram-positive bacteria, as well as the presence of appendages on the cell surface, can affect the binding with PEs.^25,26^ In particular, both groups of cells can have mechanisms of resistance to cationic PEs by incorporating molecules, which make the membrane charge more neutral.^16,25,27,28^ Cells of Gram-positive bacteria lack an outer membrane however, they have a thick cross-linked layer of a peptidoglycan.^29,30^ In contrast, cells of Gram-negative bacteria possess two cell membranes, an intermediate peptidoglycan cell wall and the outermost lipopolysaccharide layer (LPS).^31,32^ The latter creates a barrier that prevents the penetration of cationic molecules.^20,32^ Although the architecture of Gram-positive bacteria is quite uniform, the composition of the outer membrane and the LPS of Gram-negative bacteria is very diverse. Therefore, we can expect that due to this diversity, the efficiency of PE deposition onto the surface of Gram-negative cells can vary to different extents, even causing a reduced viability of cells. Although it has not been examined in past studies, Gram-negative bacteria might be a suitable study system, because the differences in the cell envelope architectures can be examined in relation to the toxicity of certain PEs.

In addition, we suppose that the duration of the cell’s exposure to PE is a factor of toxicity, which conventional toxicity tests rarely consider. The minimal inhibitory concentration (MIC) test exposes the cells to the PE present in the growth medium throughout the growth incubation period (a case we refer to as long-term or “continuous” exposure). This differs from the LBL modified system, where PEs are only deposited onto the surface of a cell forming a nano-thick capsule, while it is absent from the growth medium during incubation, because the encapsulated cells are washed by a buffer (a case we refer to as “short-time” exposure).

Therefore, we hypothesize that the toxicity of PEs depends on both physical and biological factors, i.e.: (i) the physicochemical characteristics of PE (branching, amount of the amino groups, MW, amount of PE, conditions of exposure), and (ii) the particular structure of the cell envelope. To determine the effect of coatings on the viability of bacterial cells, we selected two Gram-negative stains with different architecture of the outer layers of the well-studied *Escherichia coli* TOP10 and *Pseudomonas stutzeri* DSM 10701 based on their difference in susceptibility to positively charged PEs. Accordingly, to reveal the ambiguities in PE susceptibility we aimed: (i) to assess the interaction of the PEs and the surface of the bacterial cells, (ii) to determine the effect of the toxicity of positively charged PEs on two Gram-negative bacteria (*P. stutzeri* and *E. coli*) and (iii) to estimate changes in survivability depending on exposure conditions.

## Results

### Cationic PEs enter the *P. stutzeri* bacterial cells

In our experiments we selected two Gram-negative bacteria of *E. coli* and *P. stutzeri* to test the difference in susceptibility to positively charged PEs. Since MWs were expected to contribute to toxicity, each PE was used either in high or low MW forms. In order to find the most suitable concentration for susceptible bacteria, the experiments were performed in a gradient of PE concentrations. We also tested different conditions of cell exposure (continuous and short), since in one case, PEs can be applied as antimicrobial films, and in another they are used to make cell coatings (Table 1).

**Table 1.** Inhibition of bacterial growth estimated as the Minimal Inhibitory Concentration (MIC) for two cationic polyelectrolytes polyethylenimine (PEI) and poly(allylamine hydrochloride) (PAH) with different molecular weights. The bacterial cells of *P. stutzeri* and *E. coli* were exposed to the polyelectrolytes dissolved in the medium (continuous exposure) or they were deposited on the cell surface, washed and then resuspended in the medium (short time exposure) to determine inhibition of growth, which was quantified after 72 h.

| Polyelectrolyte | *P. stutzeri* |  | *E. coli* |  |
| --- | --- | --- | --- | --- |
|  | Continuous [µg/mL] | Short [µg/mL] | Continuous [µg/mL] | Short [µg/mL] |
| PEI 600-1000 kDa | 70.3-17.6 | 78.1-39.1 | 70.3-17.6 | not observed |
| PAH 120-200 kDa | 35.2-17.6 | 39.1-19.5 | 35.2-17.6 | not observed |
| PEI 25 kDa | 35.2-17.6 | 78.1-39.1 | 70.3-17.6 | not observed |
| PAH 17.5 kDa | 17.6-8.9 | 39.1-19.5 | 35.2-17.6 | not observed |

In the case of continuous exposure, we observed similar, statistically insignificant, toxic effects on both strains either using high or low MW PEI or high MW PAH (see Table 1). However, *E. coli* was found to be more resistant to the low MW PAH, where MIC concentrations were twice higher than in *P. stutzeri* (see Table 1). In short time exposure tests, the toxicities were strain-dependent. Accordingly, in *E. coli*, there were no toxic effects on the bacteria from both PEs either high or low MWs. We also did not observe any significant differences in the duration of the *lag* phase (data not shown) between cells exposed to the highest concentrations of the PEs and the control cells (*P* > 0.05). In *P. stutzeri* cells, we only observed higher resistance with a 2-fold increase in MIC concentrations for the low MW PAH than in continuous exposures. There were no significant differences (*P* > 0.05) in the duration of the *lag* phase (data not shown) between the PEs treated cells and the control cells.

Prior to analysis of the effect of cationic PE coatings on cell integrity, we deposited a single layer of high MW PEI on resistant cells of *E. coli* and a sensitive strain of *P. stutzeri* to observe if PEs can cause disturbance of the cell envelope (Figure 1). Although we did not see obvious disintegration of the *P. stutzeri* cell envelope, the coated cells of *P. stutzeri* had indistinct membrane boundaries compared to the coated cells. We did not observe any difference between coated and uncoated cells of *E. coli*, as they both preserved cell integrity with a clear membrane boundary.

**Figure 1.**
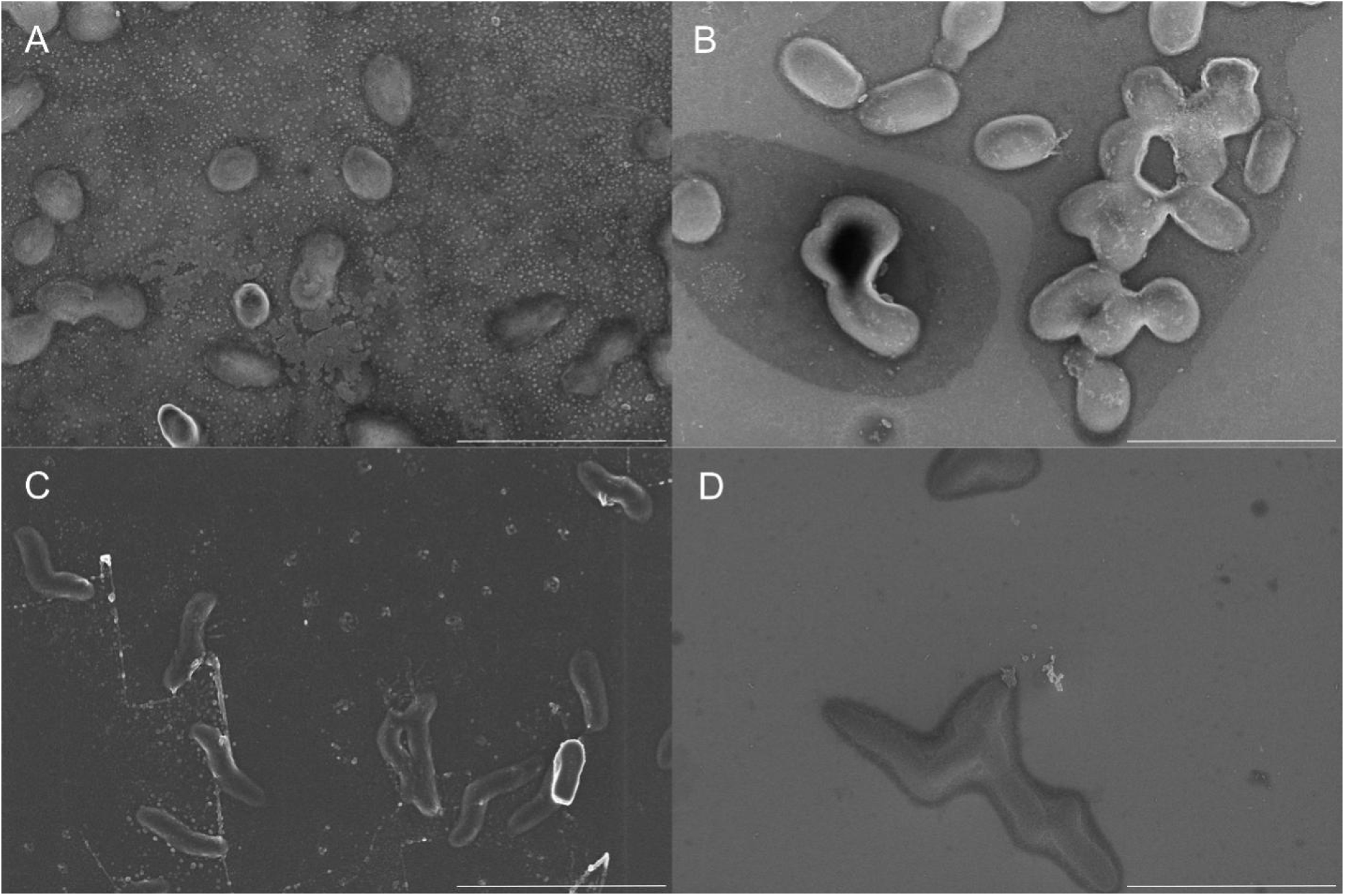
Deposition of cationic polyelectrolyte on the surface of bacterial cells. The SEM image represents uncoated cells of *E. coli* (A) and *P. stutzeri* (C) and coated cells of *E. coli* (B) and *P. stutzeri* (D) in a layer, composed of PEI 600-1000 kDa. Scale bar is 5 μm.

Since we observed a difference in susceptibility to PEs between the two strains in short time exposure, we hypothesized that PE might somehow interfere with the bacterial envelope, causing toxicity. Therefore, we assessed in detail the distributions of PEs within a single cell using super resolution STED microscopy (Figure 2). We observed that PEs penetrated the cells of *P. stutzeri* in large quantities, forming bright shining areas despite the type or MW of applied PEs. Also, the shape of *P. stutzeri* after exposure to PEs remained altered due to the formation of up to 2 bulges on bacterium at the planar plane of the membrane. Although we did not observe statistical differences (*P* > 0.1) between the normal cell diameter and the diameter of the region where the bulge was formed, the diameter with bulges was on average 1.17±0.14 larger than the diameter of the unaltered region of cells. With *E. coli*, the deposited PE layer was clearly visible on the cell surface, forming a bright intensive coating for each type of PE. Despite the signals detected inside the cells of both strains, they were on average 1.9 times lower in *E. coli* than inside *P. stutzeri* (*P* < 0.05).

**Figure 2.**
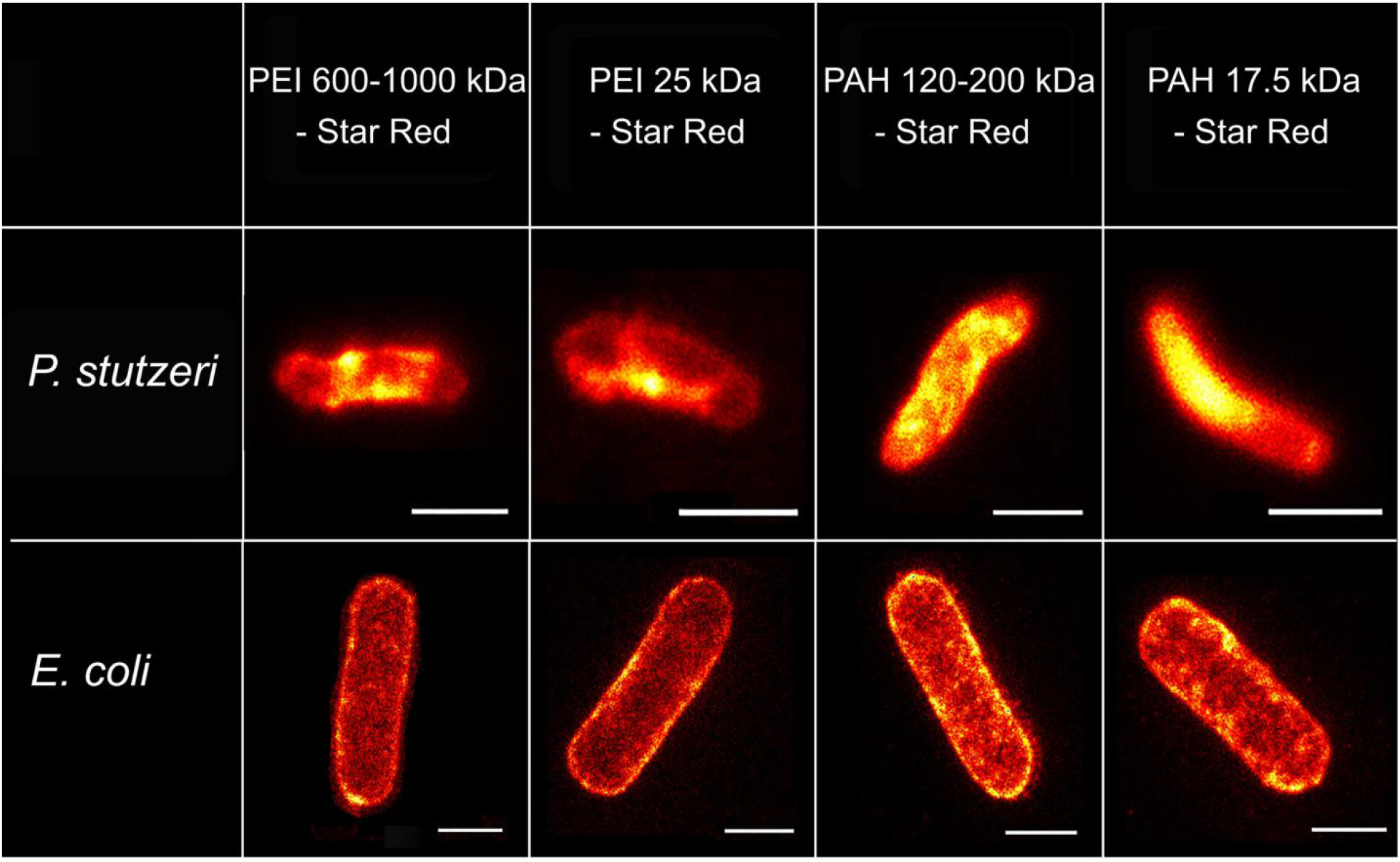
STED images of *P. stutzeri* and *E. coli* coated with polyethylenimine (PEI) and poly(allylamine hydrochloride) (PAH) polyelectrolytes with two molecular weights labelled with STAR RED STED dye. The deposited cationic PEs penetrate cells of *P. stutzeri*, while forming a capsule around *E. coli*. Scale bar for all images is 1 μm.

Analyzing external and internal signals, we determined the ratio, defining the penetration of PE inside the bacterium. The ratio was expressed as the amount of PEs on the outer layers to the amount detected inside the cells (Figure 3). In general, all PEs were distributed on the surface of *E. coli* cells (ratio > 1) and no significant changes of the ratios were observed when concentrations of PEs were increased. With *P. stutzeri* the PEs penetrated inside the bacterium (ratio < 1), and the ratio was reduced on average 1.9 times when the highest concentrations of PEs were applied (*P* < 0.02).

**Figure 3.**
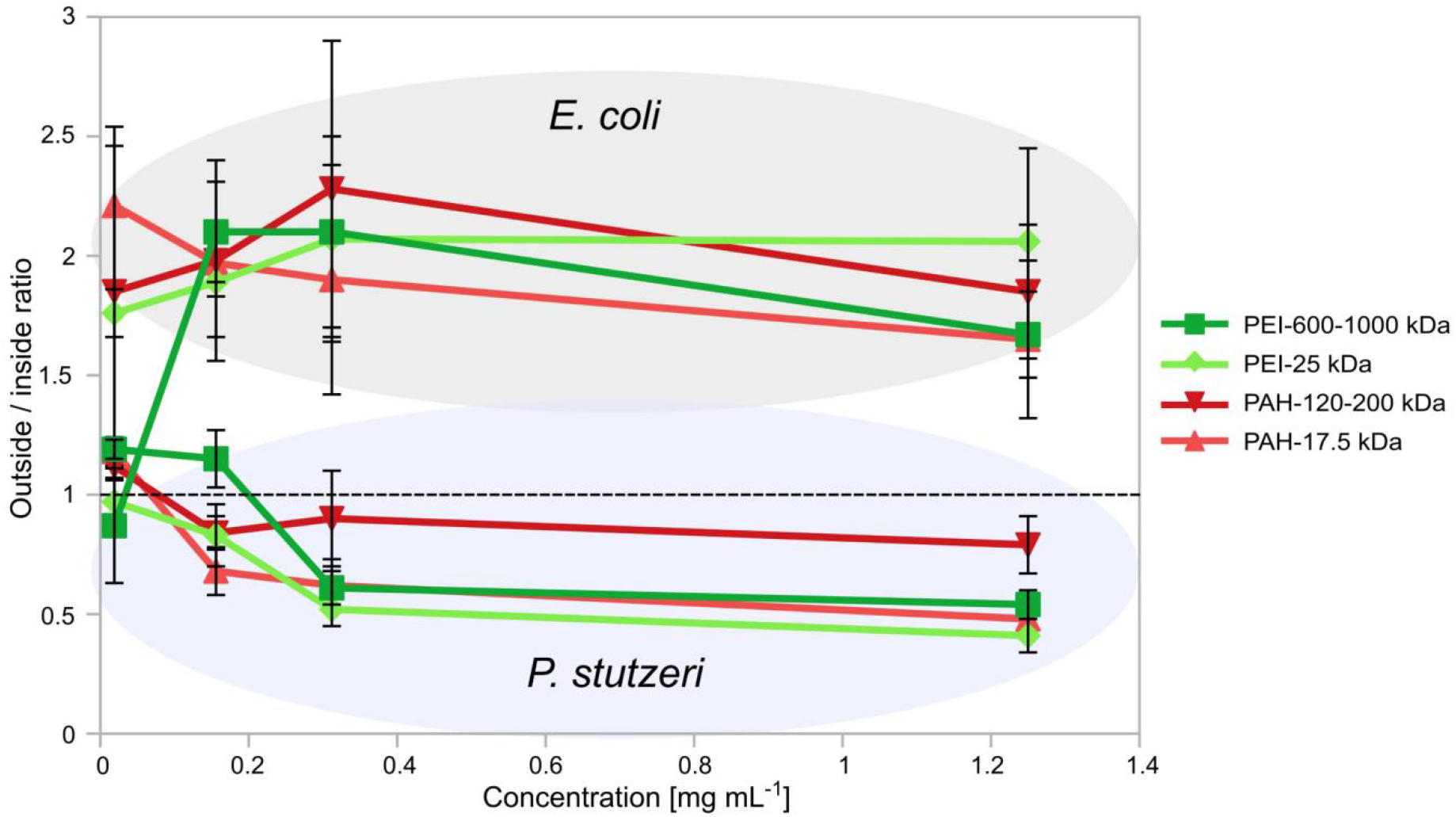
Quantitative determination of the ratio of polyelectrolytes on the outer membrane vs. the intercellular surrounding of the bacterial cells for two types of Gram-negative cells (*E. coli* – labeled in grey and *P. stutzeri* labeled in blue). The dashed line represents the boundary, above which polyelectrolytes are located on the outer layers, and below they are protruded through the cells, respectively. Two polyelectrolytes polyethylenimine (PEI) and poly(allylamine hydrochloride) (PAH) were used with two molecular weights at different concentrations. The analysis was performed on the obtained STED images.

We did not find any significant differences between the molecular weights of PEs deposited on the surface of *E. coli*, however, the intensities of both PEI were on average 2.5 times lower than PAH (P < 0.01), (Figure 4). With *P. stutzeri* the highest intensity of PEs was observed, when a low MW PAH was applied inside (358 ± 67) and outside (168 ± 14.2) the membrane. The intensities of the deposited PEs at the highest concentration were 7.8 ± 3.5 times lower on the membrane and 20.8±12.8 inside the cells of *E. coli* than *P. stutzeri* (*P* < 0.05). In addition, the correlation coefficient between the concentration and intensities of PEs for *P. stutzeri* (0.80 ± 0.12) was, on average, higher for all PEs than for *E. coli* (0.25 ± 0.26).

**Figure 4.**
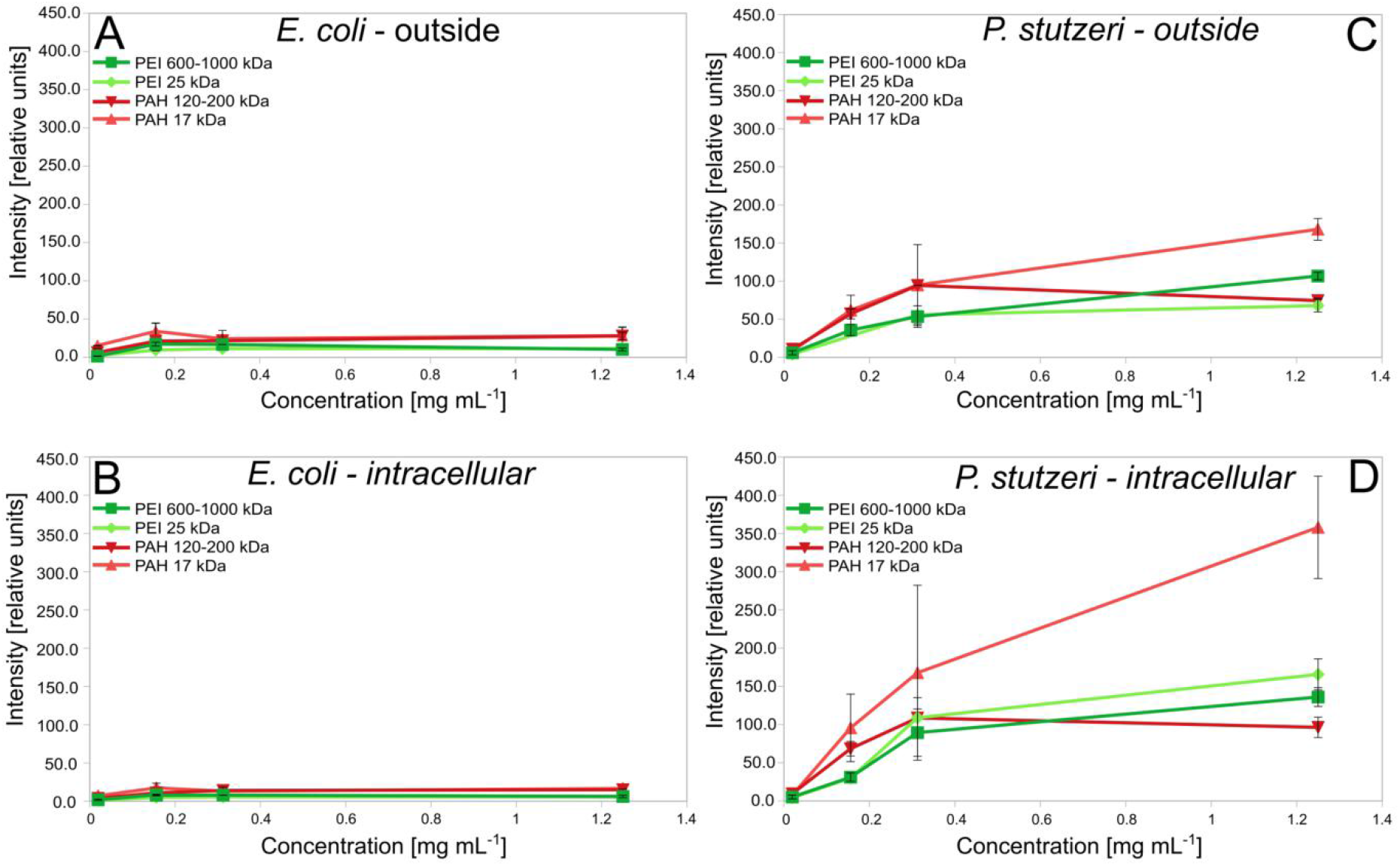
Quantitative determination of the intensities of polyethylenimine (PEI) and poly(allylamine hydrochloride) (PAH) polyelectrolytes with two molecular weights obtained from the surface or internal surrounding of bacterial cells.

### Reduction in charge densities of PEs improves viability of *Pseudomonas stutzeri*

Since the PEs were found to penetrate inside the cells of *P. stutzeri*, we tested whether there is a window, which allows to sufficiently coat the cells not being bactericidal, but keeping a positive charge suitable for further manipulations. By making serial dilutions, we estimated how the surface charge of the formed coatings depends on concentration (Figure 5A). Although the concentrations of the PEs were diluted by a factor of 2, we did not observe a two-fold decrease in the surface charge. The highest decrease of positive mobilities (1.59 times on average) from the initial concentration of PEs was only observed at 0.039 mg mL^-1^ (64 times diluted solution). The number of viable cells of *P. stutzeri* was strictly dependent on the concentration of PEs and, therefore, on the change in the electrophoretic mobility of the coated bacteria (Figure 5B). The highest reduction in Colony Forming Units (CFU) (46% ± 8% of viable cells) was at the highest concentrations of PEs (2.5 mg mL^-1^). In contrast, there was no significant reduction in growth when mobilities were negative (0.019 and 0.009 mg mL^-1^ for all PEs). Thus, in the range between the highest and lowest concentrations, we found a compensation point (above the isoelectric point) at 0.039 mg mL^-1^, where we observed the highest possible viability (73% ± 23% of viable cells) at the highest possible positive electrophoretic mobility (1.83±0.22).

**Figure 5.**
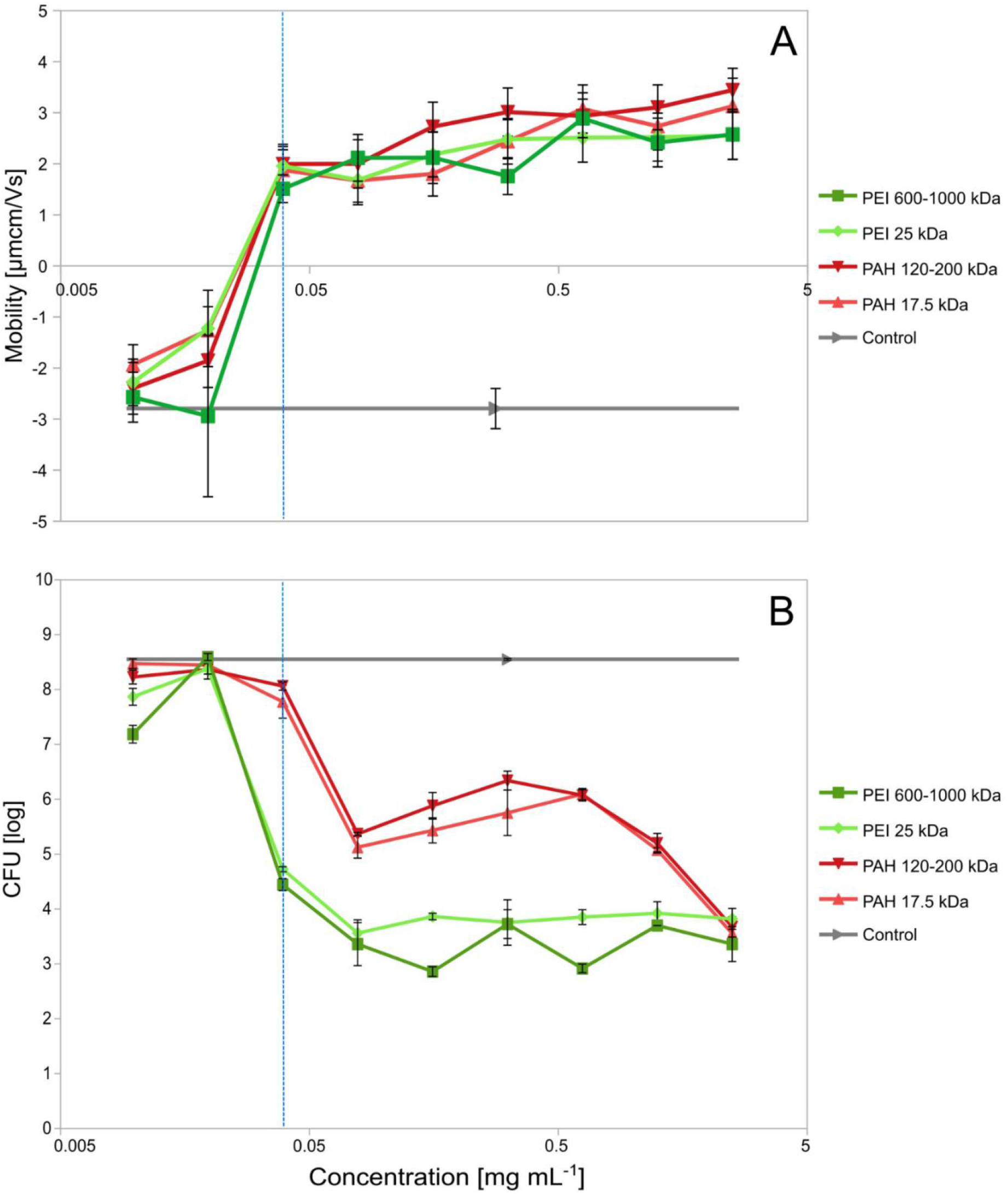
The effect of coating bacterial cells of *P. stutzeri* in a single layer of polyethylenimine (PEI) or poly(allylamine hydrochloride) (PAH) polyelectrolytes with two MWs on A) the change in electrophoretic mobility and B) viability, which was determined using plate counting of colony forming units (CFU). The dashed line indicates the compensation point, where the cells still have a positive charge and have a better viability.

The viability of the coated *P. stutzeri* cells was dependent on the particular type of PE. In general, PAH-coated cells showed significantly higher (*P* < 0.05) viabilities than PEI-coated cells between 0.625 – 0.019 mg mL^-1^. Comparisons between the same type of PEs (PEI or PAH) but of two different MWs showed no significant difference. With *E. coli*, we did not observe any significant reduction of the cell growth between control and exposed to PEs cells in all tested concentrations. Based on both experimental approaches, determining toxicities in liquid or on solid media, we observed ambiguities in susceptibility to the PEs between the tests. Accordingly, the cells were more resistant to both types of PEI in liquid surrounding, since the range of MIC concentrations was two times higher than with PAH (see Table 1). However, with using solid media, the cells were more resistant to both types of PAH than PEI (see Figure 5B).

## Discussion

Our results clearly show that PEs entered the cells of *P. stutzeri* however, they could not penetrate inside the cells of *E. coli* (Figures 2 and 3). Since we used the same PEs for two different strains, we can expect that the major contribution to permeability was related to the difference in the structure of the cell envelope. Accordingly, we summarized the specific differences in membrane composition of each bacterium as systems of protein export, LPS biosynthesis, cationic antimicrobial peptide resistance (CAMP), which can affect PE penetration (Table 2).^33–36^ The cells of *E. coli* have been found to have a much wider repertoire of mechanisms, which can help to overcome the exposure to cationic PE molecules and mask the membrane, compensation the effect of PEs. In particular, a decrease in cell-PE binding can be achieved through a decrease in the membrane charge and affinity to the PE either involving specific proteins^37^ or activating genes, which modify lipid A by ethanolamine and aminoarabinos.^27,28^ Our results also support that the membrane is somehow masked since the overall intensities of PEs were much lower in *E. coli* than in *P. stutzeri*, indicating a lower binding ability (Figure 4) and possibly, lower membrane damage (Figure 1). However, the contribution of each gene, making the cells more resistant, has to be further investigated. Although some amount of PEs was also found inside *E. coli*, it could be related either to the noise from the top or the bottom of bacterium and thus, axial limit of detection ^38^, or the cells maintain this amount of PEs inside not losing the viability. Since PEs entered the cells of *P. stutzeri*, we observed the formation of bulges (Figure 2), which can be a result of damage to the cell envelope by penetration of PEs and crosslinking of the peptidoglycan layer.^39^ In addition to morphological properties, metabolic pathways could also contribute to resistance. Both bacteria are facultative anaerobes. However, in the absence of oxygen, *E. coli* can ferment, whereas *P. stutzeri* requires nitrate. From this perspective, *E. coli* can be more resistant to hindering respiration, which can be caused by PEs.

**Table 2.**
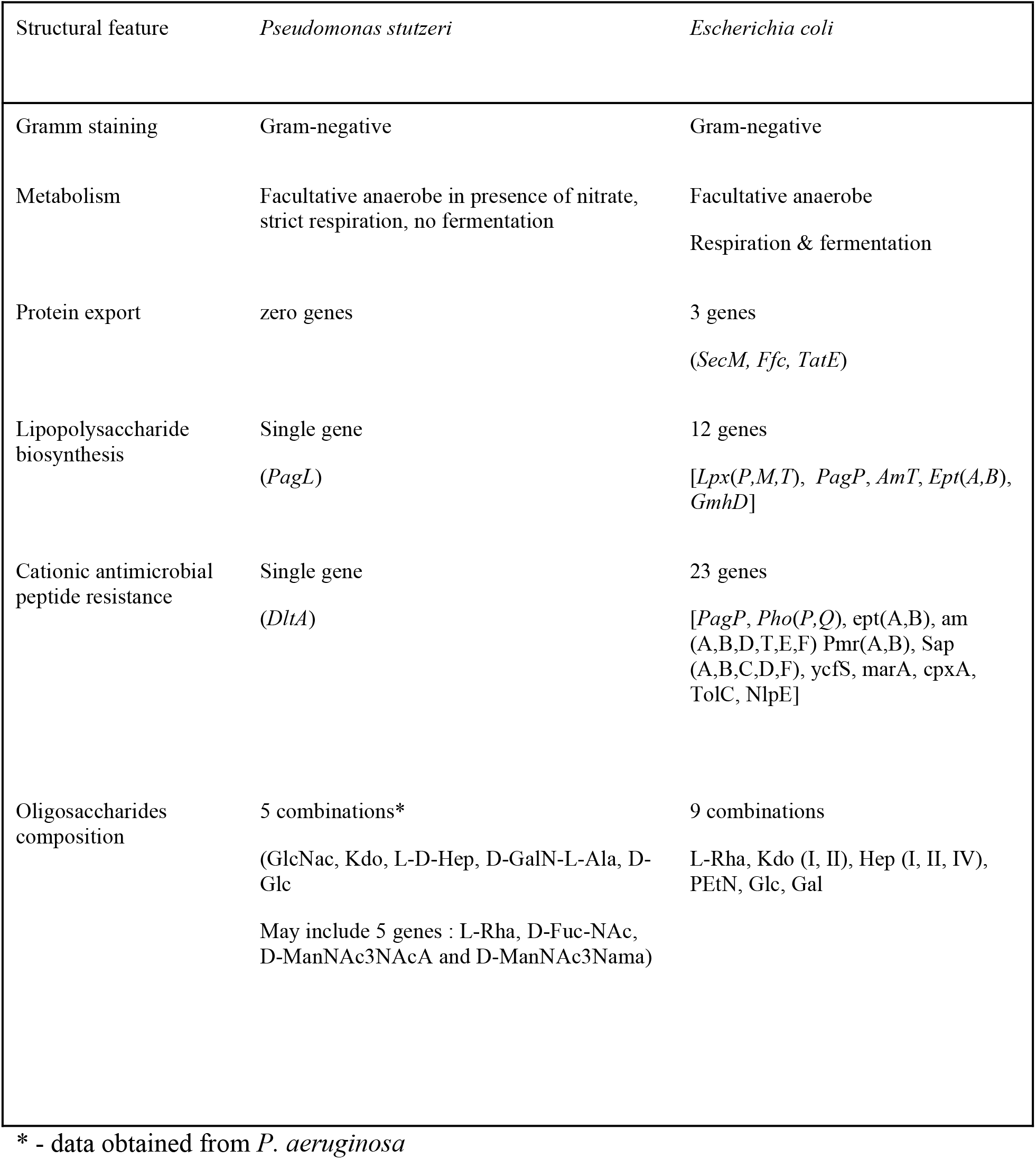
Differences in the composition and metabolism of two Gram negative bacteria. Core genes present in both strains (not shown), only unique genes for a particular bacterium are presented.

| Structural feature | *Pseudomonas stutzeri* | *Escherichia coli* |
| --- | --- | --- |
| Gramm staining | Gram-negative | Gram-negative |
| Metabolism | Facultative anaerobe in presence of nitrate, strict respiration, no fermentation | Facultative anaerobe<br>Respiration & fermentation |
| Protein export | zero genes | 3 genes<br>(*SecM*, *Ffc*, *TatE*) |
| Lipopolysaccharide biosynthesis | Single gene<br>(*PagL*) | 12 genes<br>[*Lpx(P,M,T)*, *PagP*, *AmT*, *Ept(A,B)*, *GmhD*] |
| Cationic antimicrobial peptide resistance | Single gene<br>(*DltA*) | 23 genes<br>[*PagP*, *Pho(P,Q)*, *ept(A,B)*, *am(A,B,D,T,E,F)*, *Pmr(A,B)*, *Sap(A,B,C,D,F)*, *ycfS*, *marA*, *cpxA*, *TolC*, *NlpE*] |
| Oligosaccharides composition | 5 combinations*<br>(GlcNac, Kdo, L-D-Hep, D-GalN-L-Ala, D-Glc<br>May include 5 genes : L-Rha, D-Fuc-NAc, D-ManNAc3NAcA and D-ManNAc3Nama) | 9 combinations<br>L-Rha, Kdo (I, II), Hep (I, II, IV), PEtN, Glc, Gal |

\* - data obtained from *P. aeruginosa*

The cells of *P. stutzeri*, coated with PEs at 39 μg mL^-1^, reached the compensation point at which the coated cells had positive values of mobilities (Figure 5 A) and the highest viability (Figure 5B). The number of PE molecules in this case for high MW PEI would be at least 7,3·10^4^ per single cell (at OD_600_ =0.4). However, according to our estimation, the compensation point for all PEs can be shifted even further to 32.5 μg mL^-1^ (or 6,1·10^4^ molecules per cell) in order to increase the safety of introduction of PEs to bacteria, where viability and mobility would be lg=7.93 cfu and 0.98 μmcm/Vs, respectively. By analyzing the number of CFU, it was found that PEI PEs were more toxic than PAH (Figure 5B) when the cells were incubated on solid media. PEI molecules have a larger amount of the amino groups (1N:2C atoms against 1N:3C in PAH) and higher branching, which could lead to increased binding within a thin layer of water and thus causing more damage to cells. In particular, we assume that the surface tension of a thin liquid layer around the cells on an agar plate, first, spatially arranges PE molecules to be oriented in the same plane as bacteria, and second, reduces distance between PEs and the cells, thus, a greater amount of amino groups can interact with the cell envelope. On the contrary, when cells were incubated with the PEs in a liquid medium, PAH with its primary amino groups (Table 1), exceed the quaternary counterparts in toxicity, possibly due to a more beneficial spatial orientation, which could cause stronger electrostatic forces and hydrogen bonds.^12,23^ In addition, it can be expected that excessive amount of PE could also interact with the components of the medium, reducing availability of nutrients or increasing osmolality. Accordingly, the mixture of bacteria-PE was diluted to reduce the interaction of PEs with nutrients and at to maintain the same bacteria-PE ratio in the continuous exposure test. As a result, we did not observe suppression of the growth of *E. coli*, while *P. stutzeri* still exhibited decreased viability, indicating the contribution of the constant exposure. There was also no change in the viability of *E. coli*, even by depositing multiple consecutive layers of PEs on the cell surface.^40^

The results of continuous exposure and the measured intensities on STED showed that major significant difference in PEs, which contributes to the toxicity, is the type of the amino group. The molecular weights or the amount of amino groups were not shown to contribute to the toxicity, since both PAH were not different in toxicity although they were more toxic than PEI in liquid media. In order to reduce the susceptibility of the different bacterial cells and, therefore, toxicity we propose several approaches: (i) acetylation of the amino groups in PEs, for reduction of the positive charge,^5^ (ii) the use of divalent cations, which can additionally cross-link the membrane and protect from permeabilization,^41^ (iii) genetic construction of strains, producing longer LPS tails with lowered affinity to PEs and lowered negative charge, or (iv) optimize the concentrations of PEs for keeping a small positive charge.

To summarize, LBL is an important and versatile technique for medical and biotechnological applications. Here, we showed new solutions for cases when the combination of PE and bacterium results in toxicity. Electrostatic entrapment also sheds light on differences in the membrane structure of Gram-negative cells, expanding the possibility for biomedical applications. Careful selection of PEs as well as lowering the overall concentration, can facilitate the preparation of PE coatings. Such coatings can induce expression of specific proteins (e.g. GFP), or genes responsible for quorum.^42^ Oppositely to matrix entrapment, LBL coating of an individual cell into strong porous nanometer range thick mesh enables nutrients to pass freely, but keeps cells mechanically separated from the surrounding microenvironment.^40^

## Conclusions

Our findings show new solutions for cases when the combination of PE and bacterium results in toxicity. Cell surface modification of Gram-negative bacteria has shown different outcomes in cell viability. Despite the membrane architecture being similar, structural features drastically affected the cell response. Single layer coating of *E. coli* with cationic PEs did not show any significant effects of toxicity, making them suitable templates for LbL deposition. However, *P. stutzeri* were extremely sensitive to changes in surface charges with high cell death, implying the use of careful and mild conditions for keeping the cell activity. All tested PEs could protrude through the membrane of this bacterium being detected intracellularly. We did not find any differences in toxicity with various MWs for the LbL case, however, the primary amines were more toxic in the liquid surrounding and had a higher ability to attach on the membrane based on the data from STED microscopy. In addition, we showed how variabilities in exposure affect the cell viability and how the toxicity can be reduced by manipulating the surface charge of bacteria.

## Materials and methods

### Bacteria strains and growth conditions

In our preliminary experiments (data not shown) on the coating of the cells of *Pseudomonas stutzeri* DSM10701 and *Escherichia coli* TOP10, we observed that after encapsulation, two strains exhibited opposite results in terms of viability. Therefore, we selected these strains, both from the Gram-negative group, for our experiments to determine the conditions, which make PEs toxic to one strain but not to the other. The optical density (OD_600 nm_) of each cell suspension was determined at 600 nm in a shorter light path of 200 μL well in a 96 well plate of a microplate reader (Synergy H4, Biotec). A total of 1 mL of each overnight liquid cultures (OD_600 nm_ = 1) was inoculated into a 100 mL of nutrient broth1 (NB1), (Carl Roth) and incubated by shaking at 28 °C and 37 °C, respectively, at 150 rpm until the cells reached the stationary phase (OD_600 nm_ = 1 - 1.5). Finally, the cells were collected in 50 mL tubes, centrifuged at 5000g for 5 min and washed 3 times in 30 mL sterile 0.9% NaCl. In all experiments the number of cells in suspension was normalized to OD_600 nm_ = 0.4 in a 96 well plate with 200 μL volume.

### Preparation of PE solutions for coating bacteria and determination of toxicities

Positively charged PEs with different MWs were used in toxicity experiments: poly(ethyleneimine) (PEI, MW=600 - 1000 kDa Cat.No. 03880), PEI (MW=25 kDa, Cat.No. 408727), poly(allylamine hydrochloride) (PAH, MW=17,5 kDa, Cat.No. 283215), (all obtained from Merck) and PAH (MW=120-200 kDa, Cat.No. 43092, Alfa Aesar). The PEs were suspended in MQ water (2.5 mg mL^-1^, pH = 7) and then sonicated (35kHz, 100W, 15min). Dissolved PE solutions were titrated to pH 7 using 1 M NaOH and then sterilized by filtration through a 0.2 μm sterile filter.

### Fluorescent labeling of PEs for STED imaging

Abberior STAR RED NHS carbonate was used for labeling all four PEs (PEI 600 KDa, PEI 25 KDa, PAH 120 KDa and PAH 17.5 Kda). Esters of N-hydroxysuccinimide (NHS-) react with primary amines under mild alkaline conditions (pH 7.2 to 9) and form stable amide bonds. Briefly, PEs were dissolved in MQ water at a concentration of 3.5 mg mL^-1^ and pH was adjusted to 8.5. To 3 mL of PE solution 250 mg of STAR RED dissolved in 100 μL DMSO (without water) was added under a magnetic stirring and cooled with ice. The reaction mixture was left on stirring for 18 h at 4 °C. To remove residual dye after labeling, the solution was dialyzed against purified water for 7 days using a dialysis tube with MWCO 100 kDa or MWCO 8 - 10 000 Da (SpectraPor), changing water every 12 h. After the dialyzed solution showed no traces of blue coloring, the solution was transferred into the glass vial at −40 °C and lyophilized in order to obtain dry labeled PE. Such prepared PEs were dissolved in water, pH adjusted at 7 and solutions were then diluted to obtain 0.1% (w/w).

### Deposition of PE on the cell surface

The electrostatic deposition of PEs on the cell surface was carried out as previously described.^40^ Briefly, prepared cells were exposed to one of the PE solutions with the concentration in the range of 2.5 – 0.0097 mg mL^-1^, pH 7 in MQ water, adding it to the washed cells in 1:1 v/v ratio. The highest concentration was chosen as the most suitable for coating *E. coli* in our previous study,^40^ while the lowest one was chosen because no change in electrophoretic mobilities was observed when PEs were applied to the bacteria. PEs were gradually diluted from the highest to lowest concentrations by a factor of 2 using sterile MQ solution. This suspension was incubated at room temperature (25°C) for 5 minutes. Unattached PEs were washed out from the bacterial suspensions by centrifugation at 900 g for 2 min and removing supernatant. When the PEs with lower concentrations (lower than 0.156 mg mL^-1^) were used for coating, it was more difficult to centrifuge the bacteria. Therefore, we adjusted the centrifugation speed by increasing it to 1200 g and increasing the time up to 3 min. The obtained pellet was 2 times washed by carefully adding 1 mL of 0.9% NaCl over the pellet without dissolving it. After washing, the PEs coated cells were resuspended in 0.9% NaCl solution. Both strains at the lowest concentration had a greater tendency to aggregate, so in some cases it was difficult to find a separate single cell.

### Measurement of electrophoretic mobility of PE coated cells

To determine electrophoretic mobilities the coated cells in different concentrations were washed 2 times and resuspended in 0.9% saline solution (NaCl). Prior to measurements, 50 μL of the suspension was added in 3 mL of MQ water, mixed by gentle vortexing and then measured by an ELS method (Particle analyzer, Delsa Nano, Beckman Coulter). Data were obtained from 3 replicates, where each of the measurements was produced from 15 accumulated values of the separated ELS values. In all ELS measurements the polarities of the electrodes were set fixed, with automatically adjustable voltage.

### Determination of toxicities

Toxicity was measured in two approaches, in which the cells were initially exposed to PEs by mixing the PE solution and the cell suspension in a 1:1 ratio and then (i) the PEs were either left in the cell suspension or (ii) the PEs were washed out, which we named as continuous and short exposures, respectively. Since we also preliminary observed the ambiguities in toxicities between solid and liquid media, we performed an additional test, determining colony forming units on the solid media.

### Minimal inhibitory concentration (MIC)

For testing continuous exposure, we prepared a series of two fold PE dilutions from 2.5 mg mL^-1^ to 0.0097 mg mL^-1^ by mixing 90 μL of sterile distilled water with 90 μL of PEs in a 96 well plate. Afterwards, 90 μL of twice concentrated NB was added. Finally, 20 μL of the cell suspension was added to 180 μL of a mixture of PE and medium (90 μL of PEs and 90 μL of media) and left to incubate. MIC was determined by measuring OD_600 nm_ every 30 min for 72 h by shaking and incubating at 28 °C and 37 °C for *P. stutzeri* and *E. coli* respectively with 30 s of shaking before each measurement.

Dilutions of the PEs for a short time exposure were prepared as described above. Then, diluted PE in the amount of 90 μL was mixed with an equal amount of bacteria and left for incubation for 5 min. Afterwards, the cell-PE suspension was diluted 160 times by consequent dilution of 5 μL in 195 μL of saline, shaking for 5 min, and the dilution was repeated. This dilution was chosen in order to exclude the access of the PEs from the media. A total of 20 μL of this suspension was added to 180 μL of medium. MIC was determined under the same conditions as described above. The final concentration of PE started at a PE concentration of 1.125 mg mL^-1^ and ended at 0.009 mg mL^-1^, prepared by dilution factor 2.

### Colony Forming Units (CFU)

CFU growth was assessed by plate counting at the appropriate culture dilution. The coating was performed by mixing 90 μL of PEs at different dilutions with an equal amount of bacteria and incubating for 5 min. After, all samples were diluted by mixing 10 μL of the sample and 90 μL of sterile 0.9 % NaCl for a suitable colony count. 10 μL of each dilution was transferred onto a nutrient agar plate and allowed to grow for 1 day at the appropriate growth temperature.

### Scanning electron microscopy

The coated in PEI 600-1000 kDa (2.5 mg mL^-1^) and uncoated cells (control) of both *E. coli* and *P. stutzeri* were transferred on a silicon wafer in an amount of 5 μL and left to air dry at least 2 h at room temperature (25°C). All the samples were mounted on SEM stubs and observed in a high vacuum SEM (Jeol JSM-7600F) with a field emission gun, at low voltages, to observe the structure of the cell.

### Statistical analysis

For statistical comparison two tailed t-tests (LibreOffice Calc.) were used. All experiments were performed in triplicates. Graphs were prepared and visualized using LibreOffice Calc. (The Document Foundation), and Inkscape (Free Software Foundation). Analysis of the growth curves was performed using PRECOG.^43^

### Stimulated emission depletion microscopy (STED microscopy) studies

For STED microscopy samples were prepared from a coated cell suspension, using 6 mL of cell suspension and 6 mL of Prolong Gold Antifade mountant (ThermoFisher Scientific) that were mixed on an object glass and then covered with a special STED cover glass. The samples were left for 2 h under hot air to obtain non-moving cells. The STED microscope set-up was custom build by the Abberior Instruments (lasers: 561 nm and 640 nm, STED laser: 775 nm; detectors: APD1: 560-630 nm; APD2: 630-720 nm). Image Acquisition and Analysis were performed using Imspector Software, v 0.1.^44^ High-resolution images of bacteria coated with STAR RED-labelled PE were obtained using a STED microscope with a 60x water immersion objective (NA 1.2). Image acquisition parameters were kept the same for all samples (except those with a high/low intensity) in order to ensure reliable comparison between different samples: the fluorophore was excited using an excitation laser at 640 nm (with 3 μW), its emission was depleted using a doughnut-shaped 2D STED laser at 775 nm (with 245 μW) and was detected in the range of 650-720 nm (filters by Semrock) using an APD. The dwell-time in each pixel was 30 μs, pinhole was 0.75 AU, pixel-size was set to 15 nm. The size and shape of the bacteria determined the field of view. In samples with extremely high/low amounts of PE, the excitation was accordingly decreased/increased, which was corrected in the later quantitative analysis to a laser power of the sample by multiplying or dividing to achieve the same values from a standard laser power.

A quantification of the amount and distribution of PE was programmed in Mathematica 12.0 (Wolfram Research): raw 2D STED images of bacteria were first slightly blurred using a Gaussian filter, and for each pixel the distance to the nearest background pixel was calculated using the inbuilt function “DistanceTransform” in order to obtain a list of pairs containing information on the raw intensity of each pixel and its distance to the background. The inner intensity of bacteria was calculated as the average intensity of all pixels with 2/3 dmax < di < dmax (dmax being the largest distance of any pixel from the background e.g. half of thickness of the bacteria). Edge intensity was calculated as the average intensity over a range of the edge of the bacteria. For more details, see supplementary information. Since all intensities were normalized to the laser power and derived in relative units, this allowed us to compare obtained intensities from inside and outside the bacterium as well as between two species.

## Acknowledgements

Work was supported by Slovenian-Russian bilateral project (BI-RU/16-18-039), Slovenian national projects (J4-7640, J1-6746, J3-1762, J1-9194, J7-9400, and P1-0143), Flemish-Slovenian research project: Bioavailable mercury methylation in the Adriatic sea (BE MERMAiD, grant agreement N1-0100), Helmholtz Association (grant PIE-0007 CROSSING), European Urban Initiative Actions founded project Applause (Grant agreement UIA02-228), 2019 - 2023 (EU - Horizon 2020): InteGRated systems for Effective ENvironmEntal Remediation (GREENER, grant agreement 826312) and European Commission (SurfBio project, grant no.: 952379).

## ASSOCIATED CONTENT

### Supporting Information

The procedure of quantitative assessment of polyelectrolytes deposited on the cells as well as supplementary results of the coated bacteria are present in a PDF file. The following file is available free of charge.

### Author Contributions

The manuscript was prepared through contributions of all authors. IR prepared toxicity studies and developed protocols, partially performed statistical analysis and graphs, and partially prepared the manuscript. JM performed preparation of samples for STED studies and partially performed writing. HM performed STED studies, analyzed all the data and partially performed writing, JH performed SEM analysis, MČ critically revised manuscript and elaborated ideas for SEM studies, JŠ critically revised manuscript and elaborated ideas for STED studies, JK was involved in the conceptual design and critical reading, TR prepared samples for SEM microscopy and helped to write and to structure the manuscript, AL developed the idea and concept, designed the experiments, and prepared the manuscript.

## ABBREVIATIONS

PEs: polyelectrolytes
AMR: antimicrobial resistance
MIC: minimal inhibitory concentration
PEI: polyethyleneimine
PAH: poly(allylamine) hydrochlorid
LBL: layer-by-layer
MW: molecular weight
LPS: lipopolysaccharide
CFU: Colony Forming Units

