## Supplemental methods for "Strain-dependent selective antimicrobial action of cationic polyelectrolytes on Gram-negative bacteria"

### Supplementary Materials and Methods

#### Outline of the automated procedure

#### Preparing the image for analysis

First, the 2D STED images of the distribution of StarRed-labelled polyelectrolyte on bacteria were exported from Imspector (version 16.2.8282-metadata-win64-BASE, provided by Abberior) to 16-bit tiffs. After importing the tiffs into Mathematica, an ellipse was fitted around each bacterium in order to rotate it to facilitate analysis (Figure S1). The distance of each pixel to the background (background was determined by a pre-defined threshold value) was calculated using the built-in function DistanceTransform on a Gaussian-blurred image, thus obtaining a "distance image" of the bacterium as well as pairs of values {distance, intensity} = {distance of the pixel to background on the blurred image, intensity of polyelectrolyte signal on raw image} for each pixel in the image. Because of this step, the procedure enables analysis of non-rod-shaped bacteria (objects) as well.

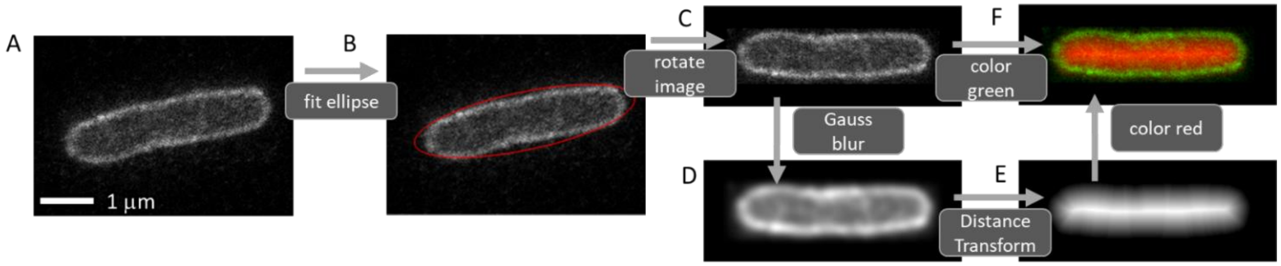


**Figure S1.** A: the original 1-channel image of StarRed-labelled polyelectrolyte on the *E. Coli* bacteria B: an ellipse is fitted around the bacteria and C: rotated. D: the image is blurred and E: the distance of each pixel to the background in the blurred image is calculated. F: For easier visualization, the overlay of the distance of each pixel to the background is shown in red (more red signal corresponds to larger distance to background) and the rotated 1-channel image is shown in green.

**Separating flat middle and curved parts of bacteria**

In order to compare the longitudinal distribution of the polyelectrolyte, the bacterium was divided into the flat middle part and curved ends of the bacteria (Figure S2). This was done by projecting the "distance image" onto the longer axis of the bacteria. The flat part of the bacterium was defined as the area between the left-most and right-most 90% of the maximal projection, the left-over regions belonging to the curved ends of the bacterium.

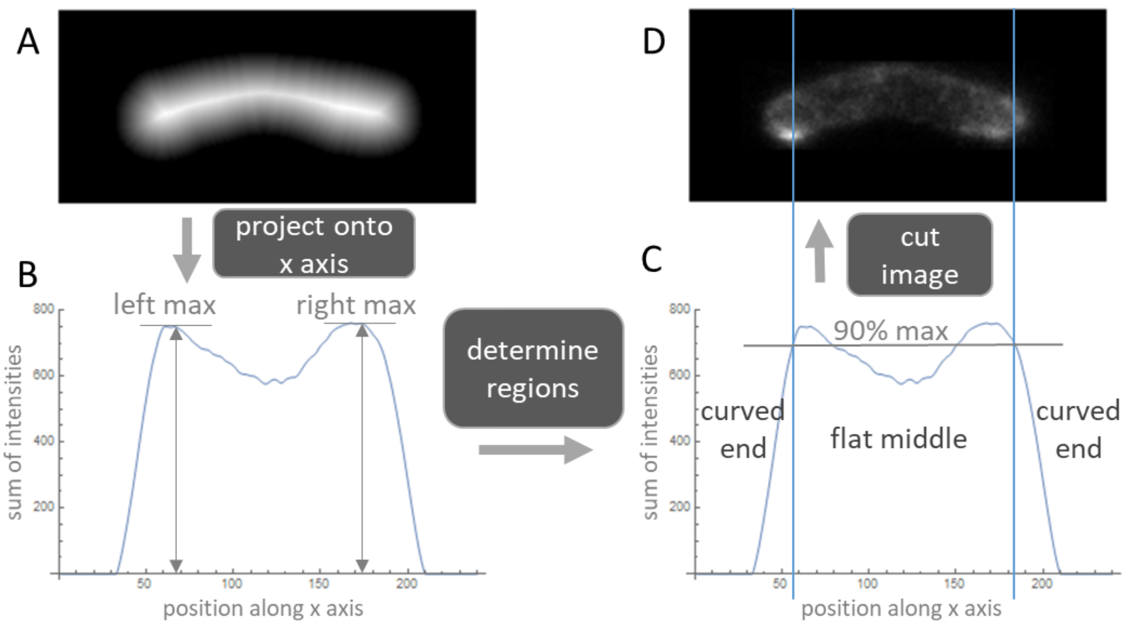


**Figure S2.** A: A distance image of the polyelectrolyte distribution on a *P. Stutzeri* bacterium is B: projected onto the major (x) axis of the bacteria. The left and right maximum are determined. C: Vertical lines at 90% of the maximal intensity of the projection indicate the boundaries between the flat and curved part of the bacteria. D: The original image of the polyelectrolyte distribution on a *P. Stutzeri* bacterium is separated into the flat and curved part of the bacterium accordingly.

**Graphs of polyelectrolyte distribution**

Next, all pairs {distance, intensity} of the bacterium image were plotted on the same graph (Figure S3). Also, two additional graphs were plotted with pixels either solely from the flat middle or from the curved ends of the bacterium. Because each pixel was plotted by a semi-transparent circle, the graphs visually convey information regarding the distribution of pixels as well.

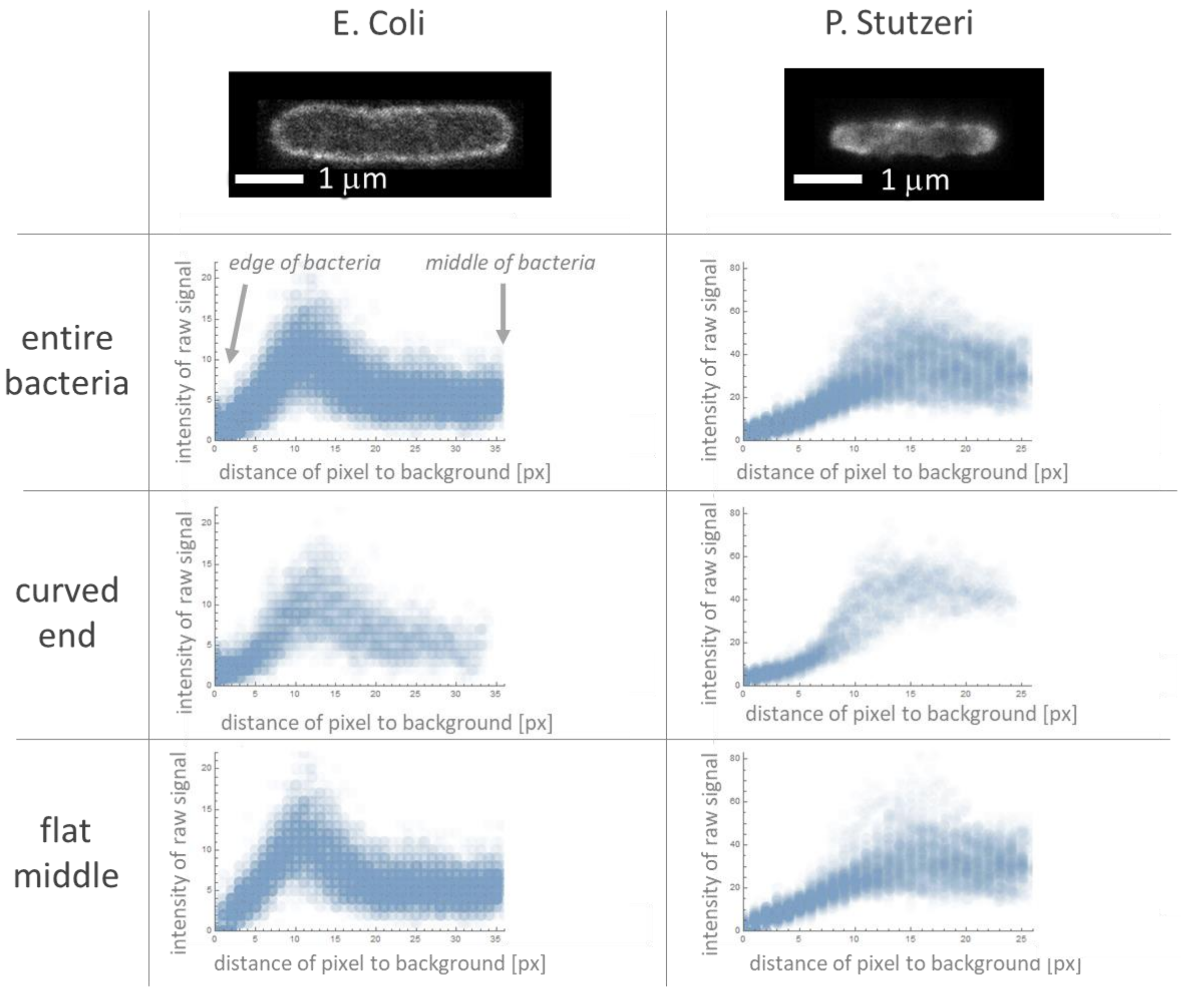


**Figure S3.** Comparison of the distribution of raw polyelectrolyte signal on two typical 2D STED images for *E. Coli* and *P. Stutzeri*.

**Quantitative assessment of polyelectrolyte distribution**

In order to obtain a quantitative assessment of the polyelectrolyte distribution for comparison with the viability tests, the data in graphs from Figure S3 was analysed. Besides the distribution of the polyelectrolyte along the longitudinal axis, we were also interested in the amount of polyelectrolyte inside and outside the bacteria. The distances of pixels to the background were used to allocate the pixel to the “interior” or “edge” of the bacteria by comparing the measured distance to pre-obtained boundary values for each region (Figure S4). For each bacteria and region (flat middle/curved end/whole bacteria, interior/edge), the mean intensity of raw signal was calculated for later comparison. The error of the mean intensity of raw signal was estimated by shifting the boundary values for interior and edge regions of the bacteria for 3 pixels – this equals to 50 nm, which is about half the thickness of the bacteria edge in *E. Coli* or 10% of the entire bacteria width.

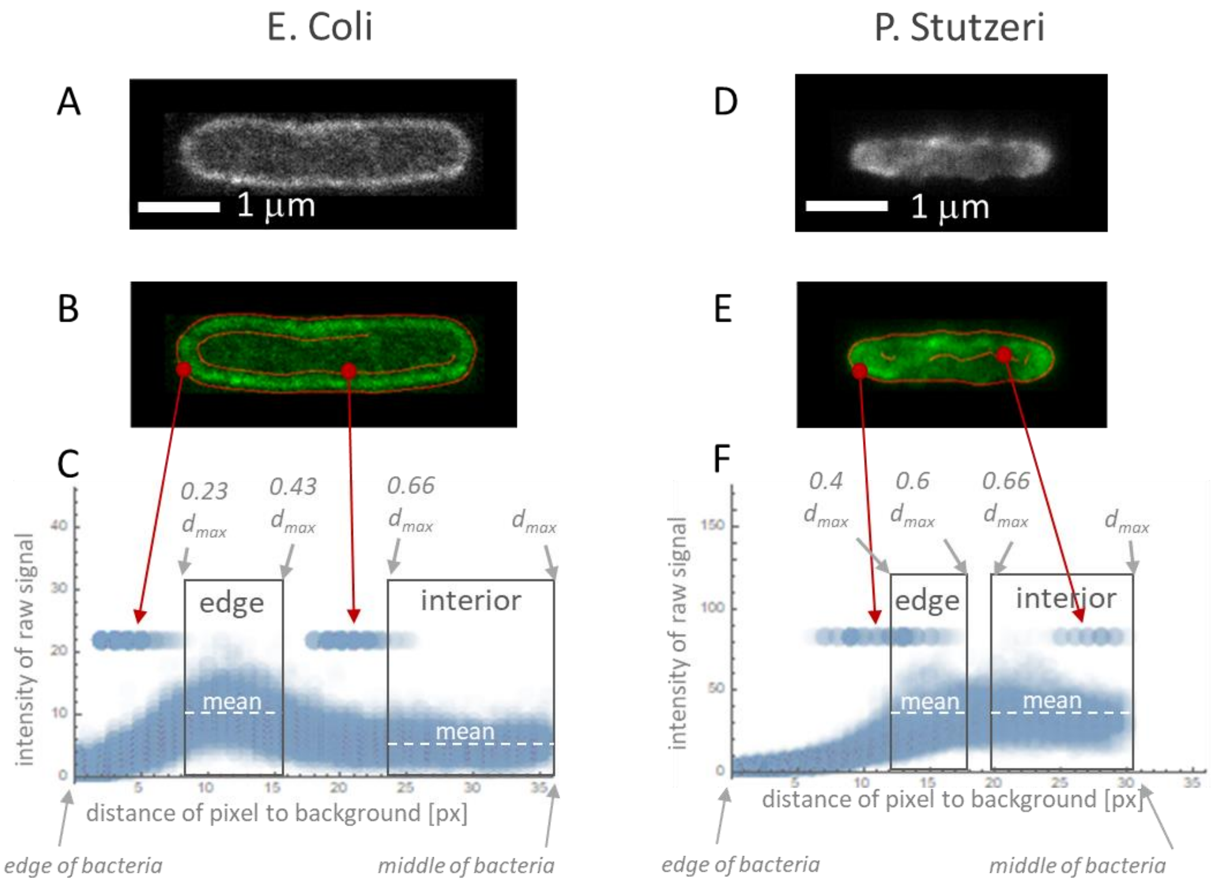


**Figure S4.** Original images of polyelectrolyte distribution for A: *E. Coli* and D: *P. Stutzeri*. B,E: Original image is shown in green and the boundaries of bacteria (obtained using the function EdgeDetect in Mathematica) is shown in red. C,F: By plotting the boundaries of bacteria from B and E on the graphs of intensity versus distance to background (same graphs as Figure S3), the boundary values of "interior" and "bacteria edge" regions for *E. Coli* (C) and *P. Stutzeri* (F) can be deducted (grey rectangles).The mean intensity in each region is denoted by a white dashed line.

**Normalisation of excitation laser power**

Since the bacterial samples exhibited a wide range of attached polyelectrolyte, they could not be imaged using the same excitation laser power. Thus, most samples were imaged using 5%

excitation, and some using 2% or 20% excitation. In order to compare these samples, the mean intensities and their errors were scaled to accommodate for the difference in excitation laser power.

**Manual analysis**

In some samples, labeled cells were aggregated, thus we did not manage to find any single bacteria – for these samples, automated analysis was not possible (*E.Coli* PEI-25 0.0019 mg/ml). To mimic the automated analysis as much as possible, we determined the interior and edge of the bacteria by eye and marked the areas in Fiji (ImageJ v1.52p) using a Freehand selection tool. The mean intensity in each region was later calculated to obtain the intensity of the polyelectrolyte in the region. To estimate the error of such a calculation, the procedure was repeated three times.

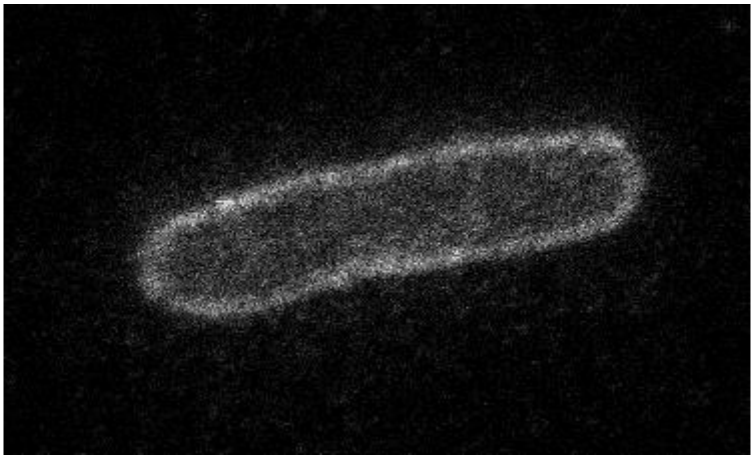

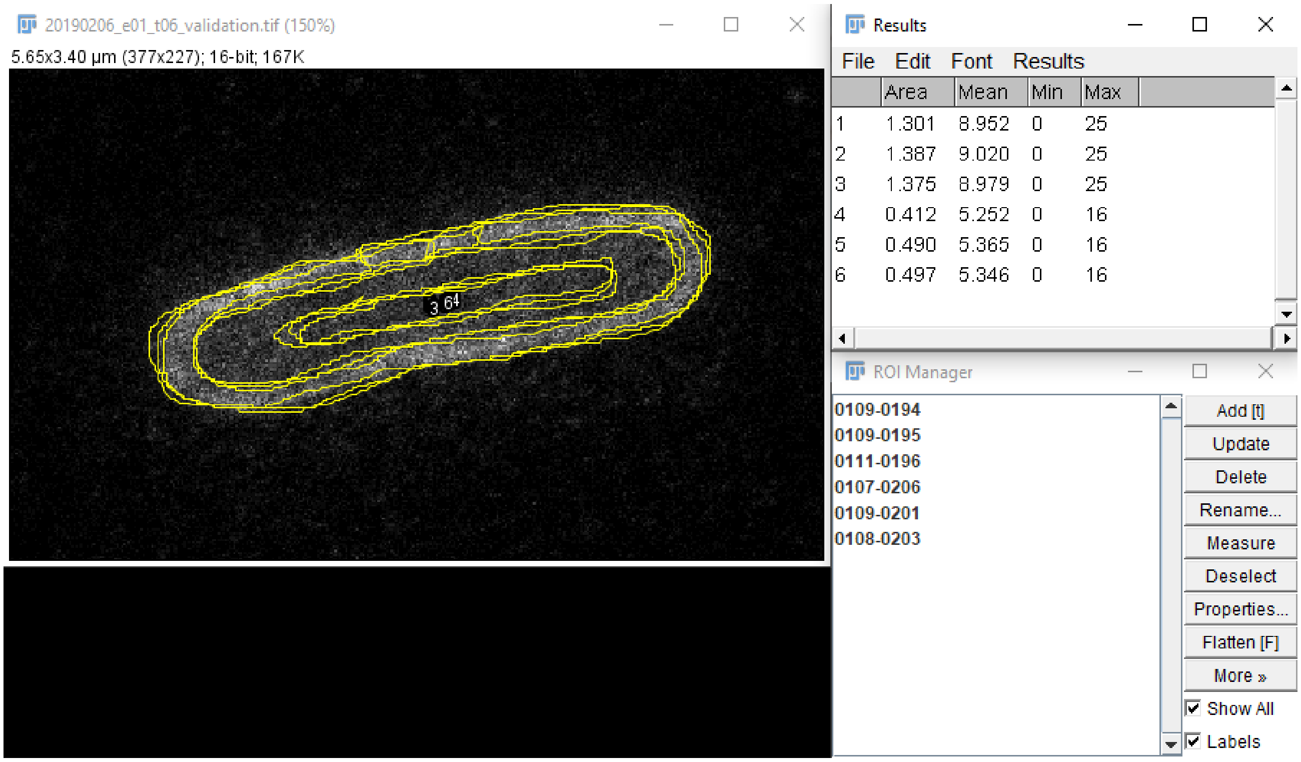
20190206_e01_t06_validation.tif (150%)
5.65x3.40 µm (377x227); 16-bit; 167K
Results
File Edit Font Results
Area Mean Min Max
1 1.301 8.952 0 25
2 1.387 9.020 0 25
3 1.375 8.979 0 25
4 0.412 5.252 0 16
5 0.490 5.365 0 16
6 0.497 5.346 0 16
ROI Manager
0109-0194
0109-0195
0111-0196
0107-0206
0109-0201
0108-0203
Add [t]
Update
Delete
Rename...
Measure
Deselect
Properties...
Flatten [F]
More »
Show All
Labels

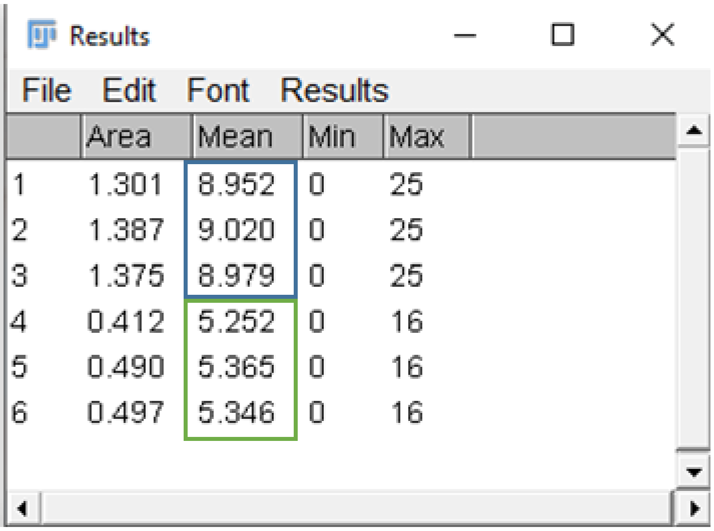

Results

File Edit Font Results

| | Area | Mean | Min | Max |
|---|---|---|---|---|
| 1 | 1.301 | 8.952 | 0 | 25 |
| 2 | 1.387 | 9.020 | 0 | 25 |
| 3 | 1.375 | 8.979 | 0 | 25 |
| 4 | 0.412 | 5.252 | 0 | 16 |
| 5 | 0.490 | 5.365 | 0 | 16 |
| 6 | 0.497 | 5.346 | 0 | 16 |

**Figure S5.** Left: Original image of the bacteria. Middle: Overlay of three repetitions of marking the “edge” and “inside” regions of the bacteria. Right: The mean intensity of the marked regions: “edge” areas 1, 2, 3 (in blue square), and “inside” areas 4, 5, 6 (in green square).

**Comparison of automated analysis and analysis by hand**

For easier comparison of the two approaches, we decided to analyse the same bacteria using both methods and compare the mean intensities on its edge and inside the bacteria. As seen in Table S1, the obtained values are in agreement with each other in the range of estimated error.

**Table S1.** A comparison of values for polyelectrolyte intensity obtained via automated and manual analysis

| | automated analysis | ImageJ – manual analysis |
|---|---|---|
| inside | 5.4 ± 0.0 (± 2) | 5.32 ± 0.04 |
| edge | 10.0 ± 1.1 (± 3) | 9.3 ± 0.2 |

### Supplementary results

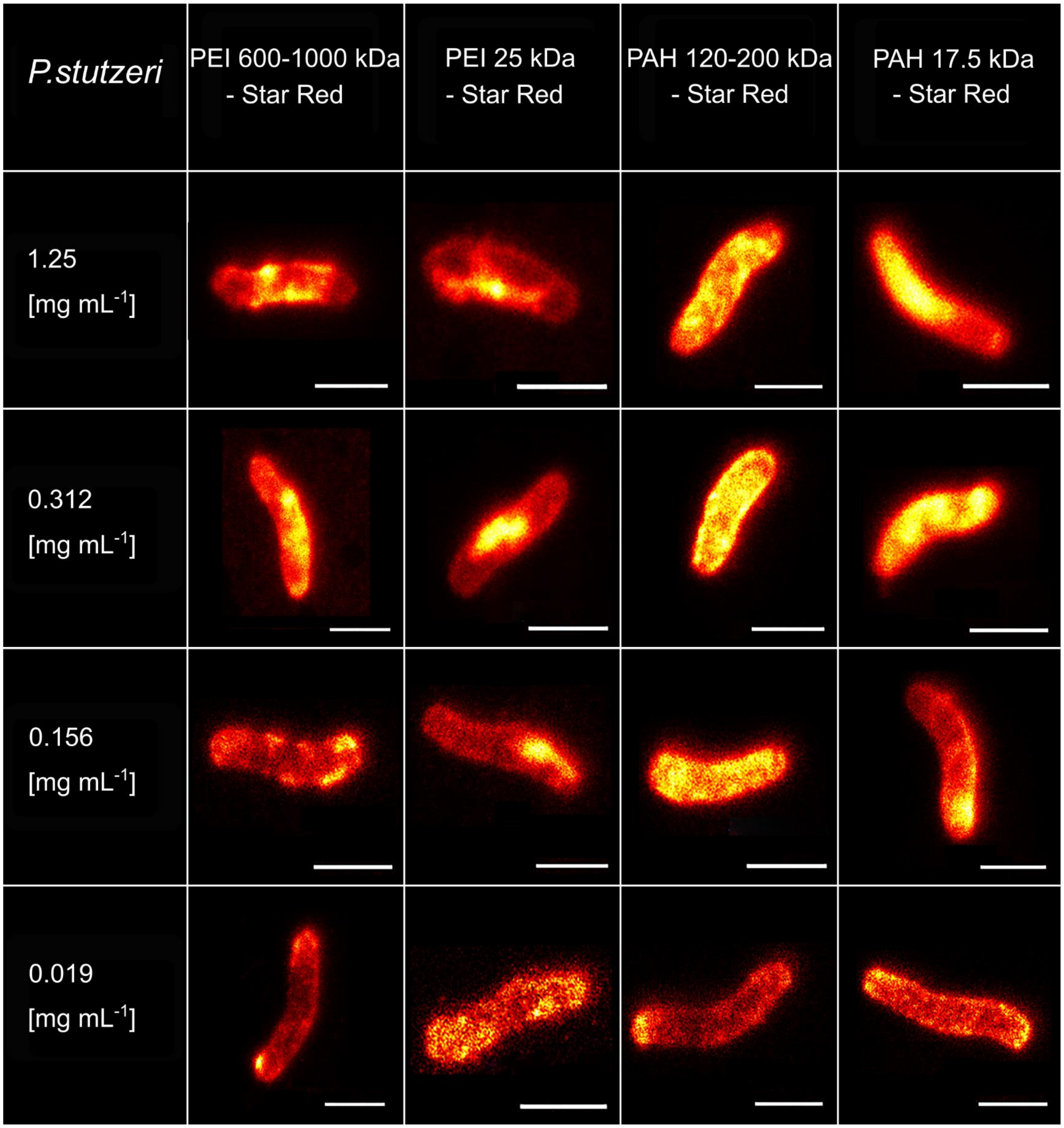


**Figure S6.** STED images of *P. stutzeri* coated with polyethylenimine (PEI) and poly(allylamine hydrochloride) (PAH) polyelectrolytes labelled with STAR RED STED dye with two MWs. Deposited cationic PEs penetrate cells of *P. stutzeri* in all concentrations. Scale bar for all images is 1 μm.

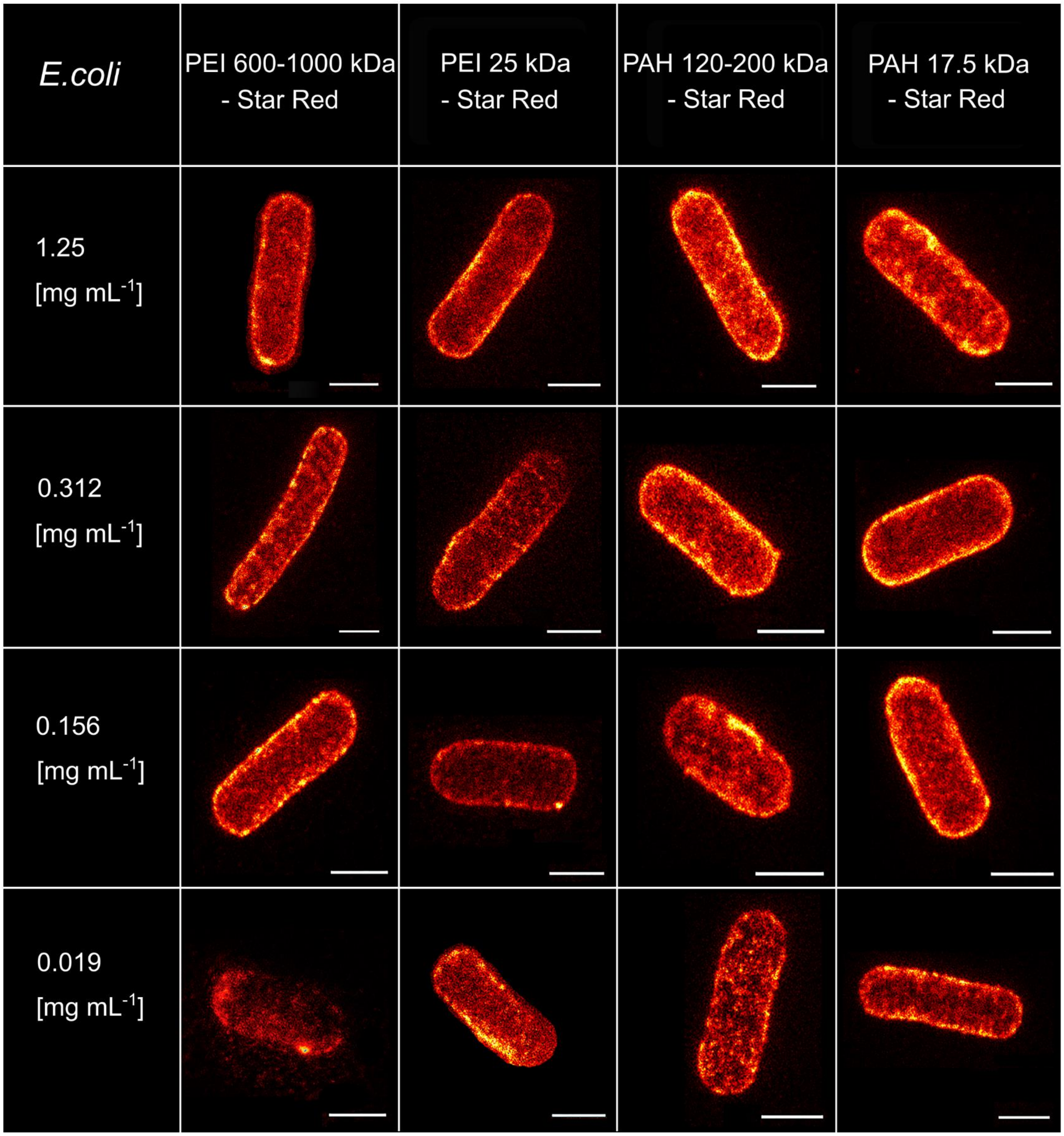


**Figure S7.** STED images of *E. coli* coated with polyethylenimine (PEI) and poly(allylamine hydrochloride) (PAH) polyelectrolytes labelled with STAR RED STED dye with two MWs. Deposited cationic PEs forme a capsule around *E. coli*. Scale bar for all images is 1 μm.
